# AI-powered pan-species computational pathology: bridging clinic and wildlife care

**DOI:** 10.1101/2022.03.05.482261

**Authors:** Khalid AbdulJabbar, Simon P. Castillo, Katherine Hughes, Hannah Davidson, Amy M. Boddy, Lisa M. Abegglen, Elizabeth P. Murchison, Trevor A. Graham, Simon Spiro, Chiara Palmieri, Yinyin Yuan

## Abstract

Cancers occur across species. Understanding what is consistent and varies across species can provide new insights into cancer initiation and evolution, with significant implications for animal welfare and wildlife conservation. We built the pan-species cancer digital pathology atlas (PANCAD) and conducted the first pan-species study of computational comparative pathology using a supervised convolutional neural network algorithm trained on human samples. The artificial intelligence algorithm achieves high accuracy in measuring immune response through single-cell classification for two transmissible cancers (canine transmissible venereal tumour, 0.94; Tasmanian devil facial tumour disease, 0.88). Furthermore, in 18 other vertebrate species (mammalia=11, reptilia=4, aves=2, and amphibia=1), accuracy (0.57-0.94) was influenced by cell morphological similarity preserved across different taxonomic groups, tumour sites, and variations in the immune compartment. A new metric, named morphospace overlap, was developed to guide veterinary pathologists towards rational deployment of this technology on new samples. This study provides the foundation and guidelines for transferring artificial intelligence technologies to veterinary pathology based on a new understanding of morphological conservation, which could vastly accelerate new developments in veterinary medicine and comparative oncology.

## Introduction

Cancers occur with phenotypically similar forms across the tree of life^1–4^. Understanding the conserved and diverged aspects of cancer across species can help answer questions about the origin and fundamental processes of its evolution. Immediate and practical advances from pan-species studies provide new tools and valuable insights into tumorigenesis and cancer resistance^5–8^, leading to improved cancer care for humans and non-human animals. Specifically, transmissible cancers presented in dogs and Tasmanian devils^9,10^ are among the few known naturally occurring clonally transmissible cancers^11^. How transmissible cancers escape immune surveillance remains unclear and is of central importance to understanding their biology and cell to cell interactions.

Despite significant resources in companion animal care, clinical treatments options are limited for a few aggressive cancers in dogs^12,13^ that represent one of the best models of human cancer^14^. Beyond domesticated species, various studies have identified valuable models in wildlife^15^. For instance, the naturally-emerging urogenital carcinoma in California sea lions^16^ and papillomavirus triggering brain tumours in raccoons^15^ are remarkable examples of pathogen-driven neoplasms. Animals managed in zoological institutes also exhibit occurrence of neoplastic growth according to several international studies, including, a 10-year survey in the Taipei zoo, Taiwan^17^, a study of cancer development in vertebrates in French zoological parks^18^, a 42-years of mammals necropsy data compilation from the San Diego Zoo, United States^19^, and a report on renal lesions followed by neoplastic and inflammatory responses in captive wild felids in Germany^20^. Studies of these animals can provide unique insights into the biology and evolution of cancer across the tree of life towards improving animal welfare by early detection and helping conserve endangered species^21,22^.

Challenges for establishing a unified comparative oncology agenda include sample collection, data management, analysis, and integration^23–27^. These can be tackled by incorporating artificial intelligence (AI) algorithms, which can empower veterinary pathology and help dissect the complexity of cancer across species and scales, from genes to epidemiology. Computational pathology powered by AI has revolutionised the study of human cancers and helped improve our understanding of the immune microenvironment^28^. In contrast to human cancer management, we lack systematic and standardised AI protocols and digital archiving and analysis of samples to study animal cancers; hence, veterinary research has not fully adopted digital pathology^25^ although efforts are being made to move forward internationally adopted guidelines for tumour pathology^27^.

Hence, we propose AI has the power to fuel pan-species tumour histology and efficiently manage data-related bottlenecks. Thus far, computational pathology in the study of non-human cancers, and non-human pathology in general, is very limited^24,25^. Convolutional neural networks have been applied to detect mitotic activity from histological slides of canine cancers^13,29^. In sheep, deep learning has been employed to delineate growth phases of mammary development^30^. Other machine learning techniques have been used to classify a common gastrointestinal disease in cats^31^. Along with computational pathology, incorporating AI into the veterinary practice of imaging techniques such as CT scans, magnetic resonance imaging, and positron emission tomography^32^ encourages the development of integrative clinical care. Such an integrative approach promises to direct precision medicine in veterinary oncology by tailoring strategies for individual patients. It includes classifying patients who differ in their treatment response and/or prognostic outcomes.

In this work, we explore and exploit the conservatism of cell morphology in neoplasias across species by applying an AI tool trained in human lung cancer^33^ (Fig 1). We evaluate the accuracy of this AI tool in mapping tumour cells distribution and lymphocytic infiltration in histological tissues from transmissible cancers and its generalisability to 18 other species. To the best of our knowledge, this is the first effort to apply computational pathology algorithms to transmissible cancers and pan-species pathology beyond mammals, thereby decoding the composition of cells in tumours across species. Our approach aims to pave the way for pan-species comparative pathology and contribute to understanding the emergence and prevalence of cancer in nature.

**Figure 1.**
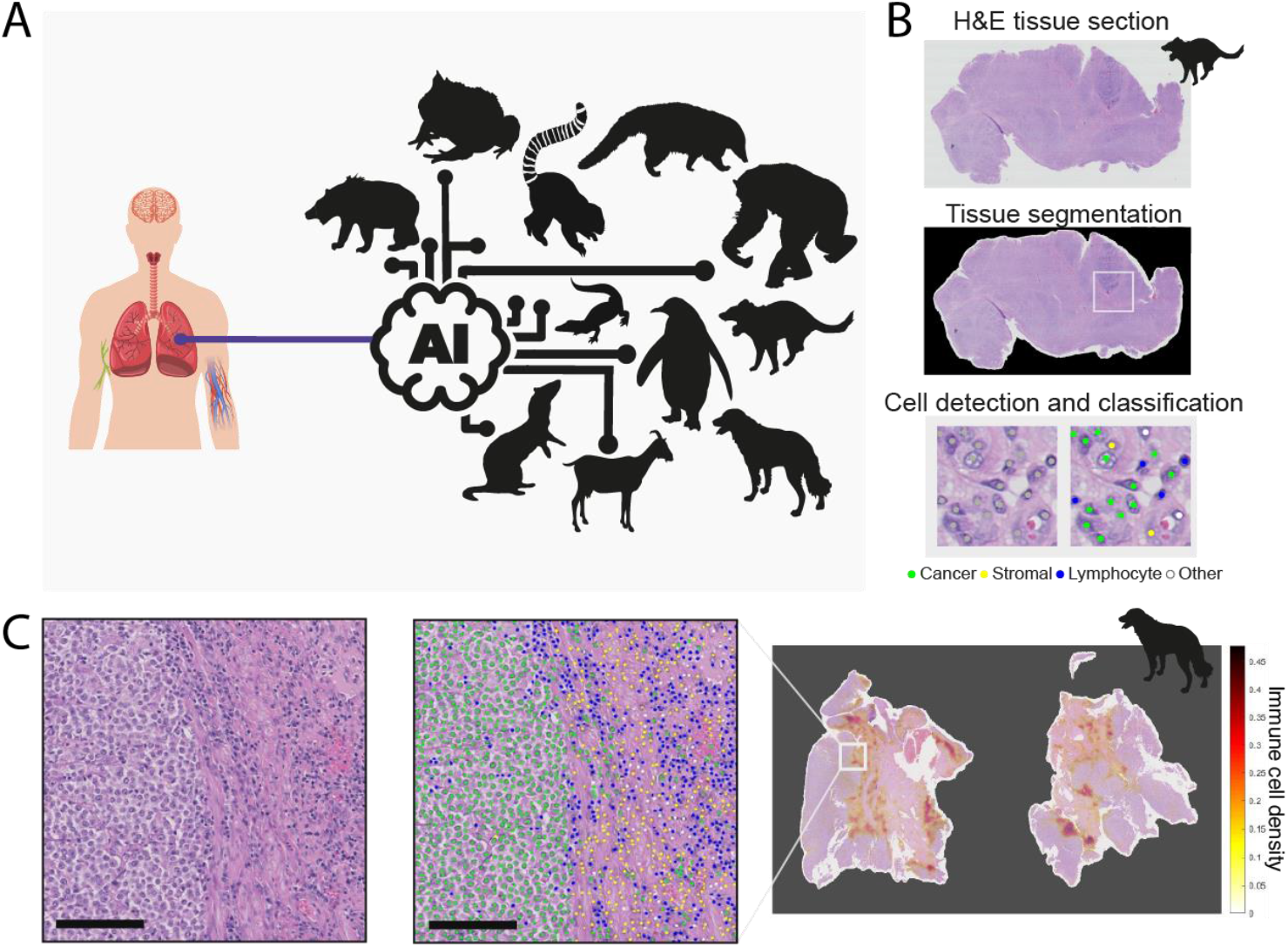
Pan-species computational pathology. (A) *Transfer learning* of cell identification from human lung to pan-species tumour pathology. (B) Overview of the H&E single-cell analysis pipeline illustrated from a Tasmanian devil’s (SARHAR) facial tumour. This AI pipeline ^33^ first segments the viable tissue area, then detects and classifies all cells into cancer, stromal, lymphocyte and others. For more details, see Methods. (C) The same pipeline is implemented to spatially profile the immune microenvironment in a dog’s (CANFAM) transmissible venereal tumour. Scale bar, 250 μm. Cell colours are denoted as four training classes, green: cancer (malignant epithelial) cells; blue: lymphocytes (including plasma cells); yellow: noninflammatory stromal cells (fibroblasts and endothelial cells); white: ‘other’ cell class that included nonidentifiable cells, less abundant cells such as macrophages and chondrocytes and ‘normal’ pneumocytes.

## Results

### Collection and quality control for veterinary histology samples

Ten hematoxylin and eosin (H&E)-stained tumour samples from 3 individuals with Tasmanian devil facial tumour disease 1 and 2 (DFT1 and DFT2) and 6 with canine transmissible venereal tumour (CTVT) were collected and digitalised from the Transmissible Cancers Group, University of Cambridge. Of these, 7 passed visual quality control for image analysis. One representative slide was chosen by the pathologists for each species considering scanning resolution and level of immune infiltration in the tumour microenvironment. In addition, H&E samples from 18 species were selected from the Zoological Society of London’s (ZSL) pathological archive and digitalised (classes Mammalia = 11 species, Reptilia = 4, Aves = 2, and Amphibia = 1). The neoplastic lesions were broadly categorised into five main tumour groups: round-cell (n = 4), epithelial (n = 9), mesenchymal (n = 4), neuroendocrine (n=2) and sex-cord stromal (n=1) tumours. A rich, pan-species digital pathology atlas was created, providing digital slide images, digitalisation and quality control protocols, and pathological annotations described below.

### Transferring AI technologies to non-human species

A deep learning pipeline tailored for human lung cancer (predominantly lung adenocarcinoma, including lung squamous cell carcinoma^33^, Fig. 1A) was applied without modification to all 20 H&E samples. Briefly, this pipeline identifies the precise location of individual cells in each H&E and classifies them based on nuclear morphology in one of four cell types: tumour cells, lymphocytes, stromal cells (fibroblasts and endothelial cells) and ‘other’ cells (macrophages, pneumocytes and non-identifiable cells) (Fig. 1B-C). We evaluated the accuracy of the convolutional neural network (CNN) with 14,570 cancer, lymphocyte, and stromal single-cell annotations from two board-certified specialist veterinary pathologists (CP and KH). For each slide, we computed the algorithm’s balanced single-cell classification accuracy (BCAcc, Table 1), as well as F1 score, precision, sensitivity and specificity (Figs S1–S2).

**Table 1.** Summary of overall balanced classification accuracy (BCAcc) by species. Balanced accuracy is computed as the average of sensitivity and specificity, ‘overall’ refers to the average of cancer, stromal and lymphocyte cells.

| Code | Common name | Species | Diagnosis | Neoplasia site | Tumour type | Annotations | BCAcc |
| --- | --- | --- | --- | --- | --- | --- | --- |
| BITARI | Puff adder | *Bitis arietans* | Carcinoma | Pancreas | Epithelial | 336 | 0.88 |
| CANFAM | Dog | *Canis familiaris* | Canine transmissible venereal tumor | Intra vaginal | Round-cell | 629 | 0.94 |
| CAPHIR | West African pygmy goat | *Capra hircus* | Lymphosarcoma | Forestomach | Round-cell | 965 | 0.70 |
| CRAHEA | Panay cloudrunner | *Crateromys heaneyi* | Hepatocellular carcinoma | Liver | Epithelial | 730 | 0.89 |
| CYACYA | Red-legged honeycreeper | *Cyanerpes cyaneus* | Sertoli cell tumor | Testis | Sex-cord stromal | 762 | 0.86 |
| DASBYR | Kowari | *Dasyuroides byrnie* | Squamous cell carcinoma | Mouth | Epithelial | 462 | 0.74 |
| GALMOH | Greater bushbaby | *Galago moholi* | Squamous cell carcinoma | Skin | Epithelial | 684 | 0.79 |
| GONOXY | Redtailed ratsnake | *Gonyosoma oxycephala* | Metastatic anaplastic sarcoma | Multiple | Mesenchymal | 526 | 0.91 |
| LEMCAT | Ring-tailed lemur | *Lemur catta* | Haemangiosarcoma | Kidney | Mesenchymal | 1049 | 0.79 |
| LEOCHR | Golden-headed Lion Tamarin | *Leontopithecus chrysomelas* | Adenoma | Pituitary | Epithelial | 601 | 0.94 |
| LEPFAL | Mountain chicken frog | *Leptodactylus fallax* | Adenocarcinoma | Celomic cavity | Epithelial | 740 | 0.81 |
| MELURS | Sri Lankan sloth bear | *Melursus ursinus inornatus* | Pheochromocytoma | Adrenal | Neuroendocrine | 959 | 0.88 |
| MUSPUT | Domestic polecat | *Mustela putorius furo* | Sebaceous epithelioma | Skin | Epithelial | 702 | 0.88 |
| NASNAS | Brown-nosed coati | *Nasua nasua* | Lymphosarcoma | Multiple | Round-cell | 520 | 0.57 |
| OSTTET | West African dwarf crocodile | *Osteolaemus tetraspis tetraspis* | Lipoma | Liver | Neuroendocrine | 1142 | 0.77 |
| PANTRO | Chimpanzee | *Pan troglodytes* | Spindle cell tumor | Palate | Mesenchymal | 866 | 0.75 |
| SARHAR | Tasmanian devil | *Sarcophilus harrisii* | Devil facial tumor 1 (DFT1) | Hard palate near left side | Round-cell | 484 | 0.88 |
| SPHHUM | Humbolt penguin | *Spheniscus humboldti* | Renal cell adenoma | Kidney | Epithelial | 452 | 0.72 |
| SUSBAR | Bearded Pig | *Sus barbatus* | Adenocarcinoma | Uterus | Epithelial | 1595 | 0.80 |
| VARPRA | Emerald monitor | *Varanus prasinus* | Spindle cell sarcoma | Multiple | Mesenchymal | 366 | 0.80 |

For evaluating the accuracy in classifying cells by the algorithm, we compared its predictions against veterinary pathologists’ annotations. The algorithm’s average balanced accuracy across cell classes showed a diverse range of variation between and within tumour groups (Figs. 2A, S1–S2). Tumour types have the same overall accuracy for cell classification based on the balanced classification accuracy values (LR test, overall BCAcc averaged across samples = 0.81; LR test, χ^2^[3] = 0.314, p = 0.957). Moreover, despite the heterogeneous number of annotations per tumour type (Fig. 2B), the balanced accuracy was not associated with the number of annotations (Spearman’s ρ = 0.088, p = 0.71) (Fig. 2B-C).

**Figure 2.**
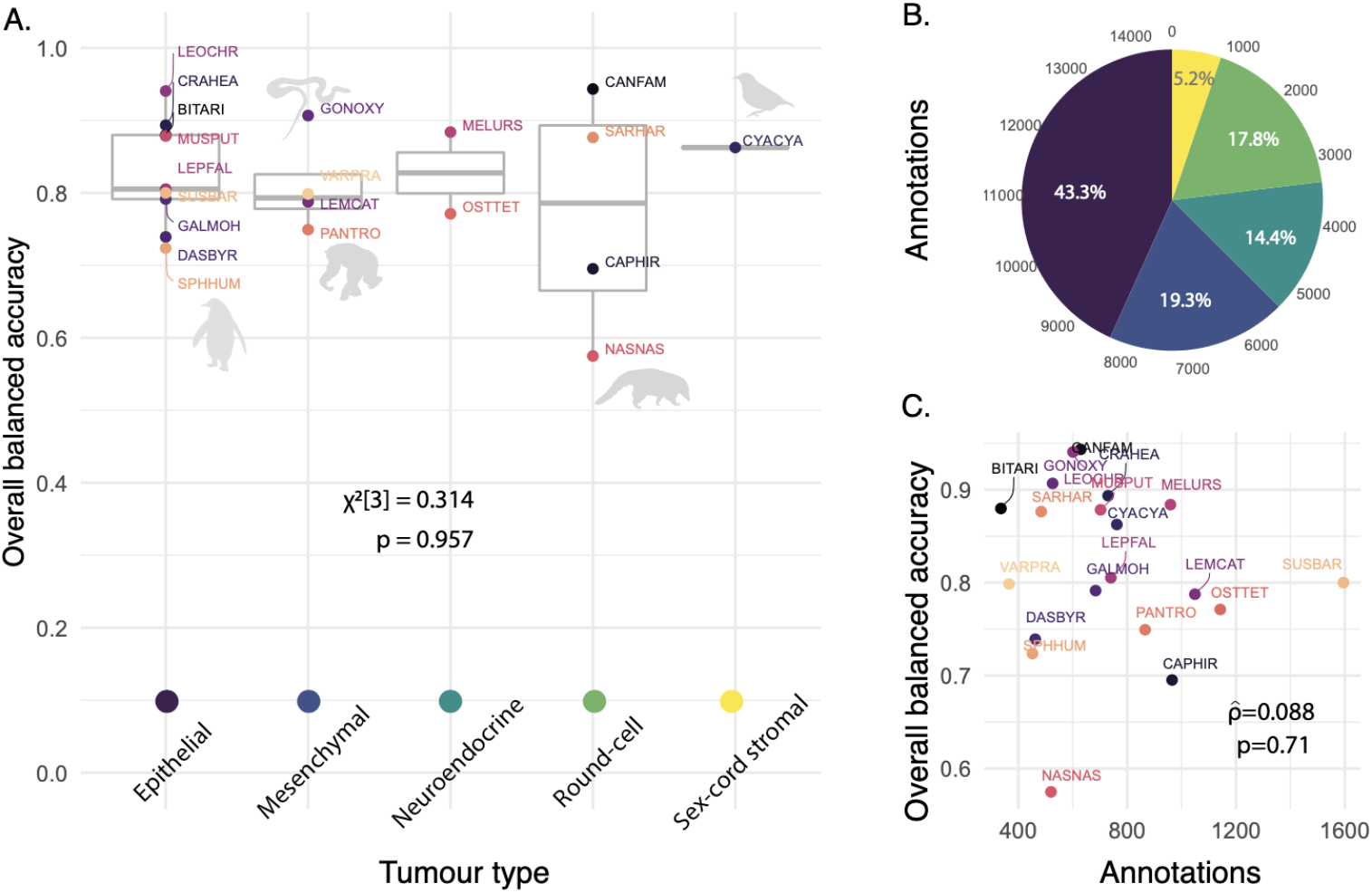
AI single-cell prediction comparison across tumour types. Balanced accuracy is computed as the average of sensitivity and specificity, ‘overall’ refers to the average of cancer, stromal and lymphocyte cells. (A) Pan-species overall balanced accuracy grouped by tumour type. (B) Distribution of the number of annotations by tumour type (colours correspond to tumour type in A). (C) Relationship between the number of annotations and the overall balanced accuracy for each species using Spearman’s correlation. Species in (A) and (C) are labelled with their codes, for more species information, see Table 1.

### Consistent accuracy across tumour types but higher in mammals

Overall, the model’s best performance was mainly in mammals (Fig. 3). In particular, the AI algorithm achieves high accuracy in measuring immune response for the two transmissible cancers (canine transmissible venereal tumour - CTVT, 0.94; Tasmanian devil facial tumour disease-DFTD, 0.88). The canine transmissible venereal tumour (in *Canis l. familiaris*) exhibited the best accuracy across all 20 species (overall precision = 0.98, F1 and BCAcc = 0.94, Fig. 3). Surprisingly, in the metastatic sarcoma in a snake (*Gonyosoma oxycephalum*), the CNN also reached a high accuracy (Fig. 4A, overall precision = 0.89, F1 = 0.89 and BCAcc = 0.91).

**Figure 3.**
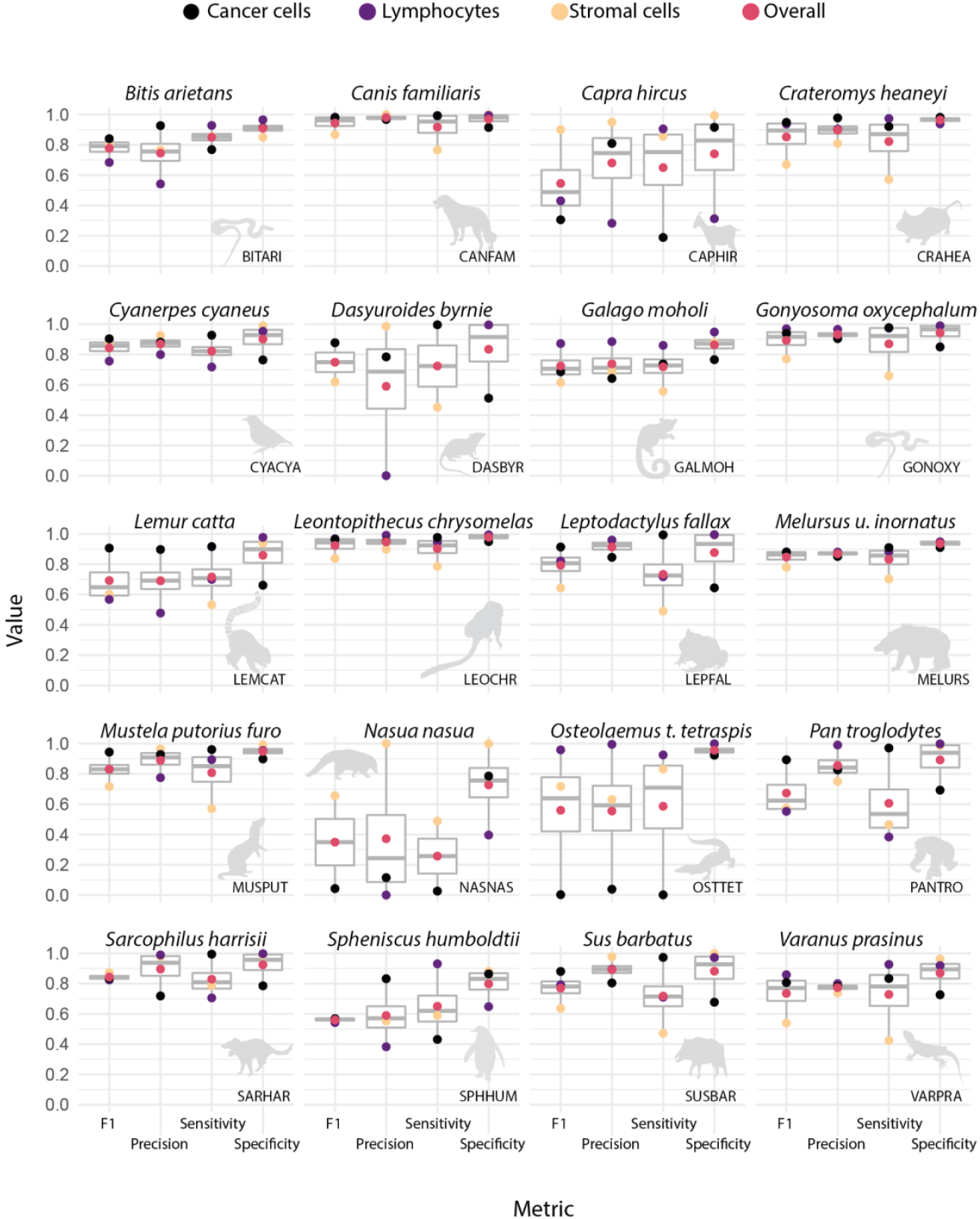
AI prediction variability for inter and intra-species tumour microenvironment cells. For each species, four metrics were evaluated including F1, precision, sensitivity and specificity (as labelled on the bottom x-axis) for the prediction accuracy of cancer, lymphocyte and stromal cells as well as their average shown as ‘overall’ (as denoted with colours on the top x-axis). For species codes, see Table 1.

**Figure 4.**
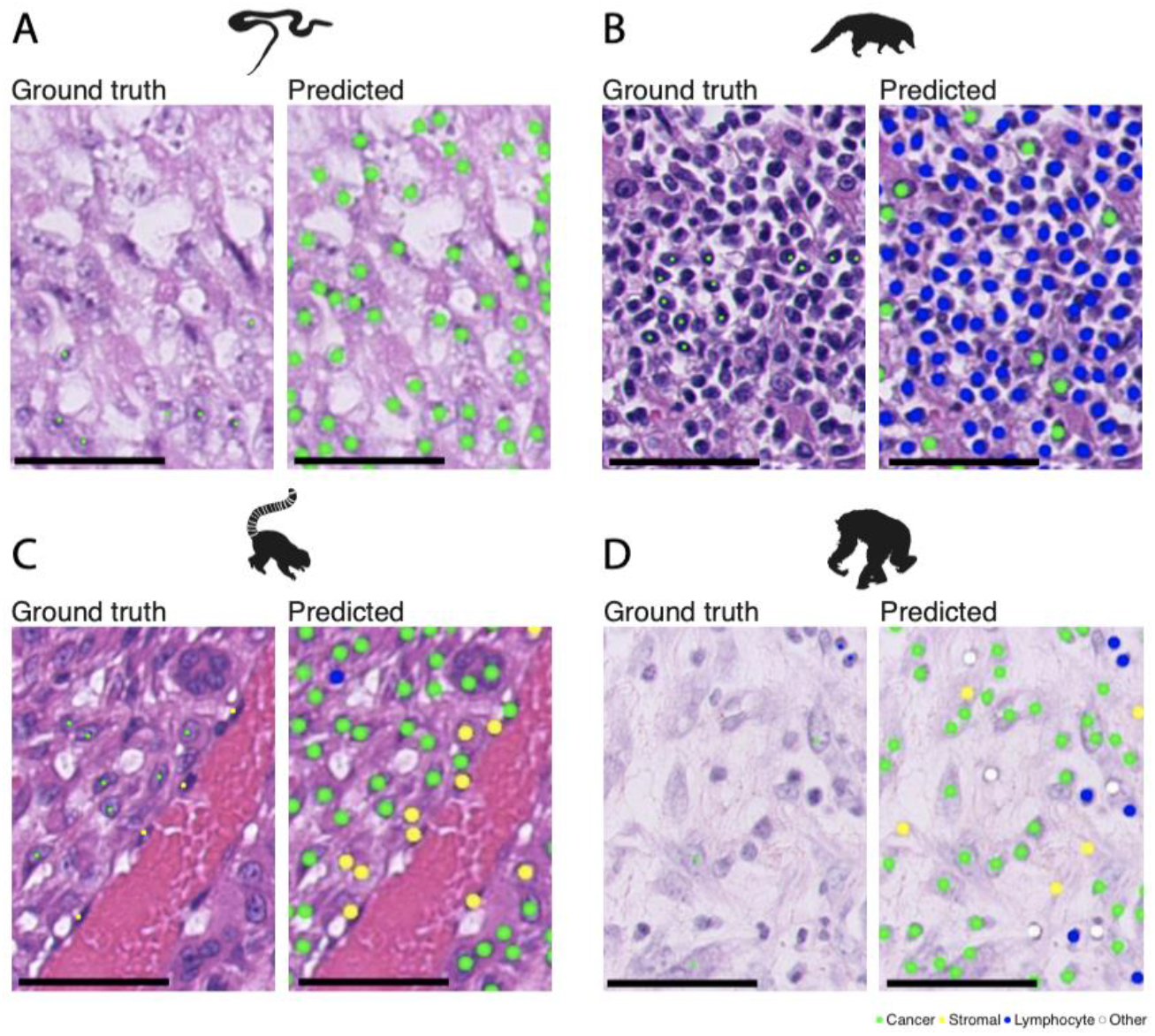
Strengths and pitfalls of current methods. Each H&E example is shown as a raw image with expert pathology annotations on some cells (left) and AI cell identification (right). Scale bar, 100 μm. Cell colours are denoted as four training classes, green: cancer (malignant epithelial) cells; blue: lymphocytes (including plasma cells); yellow: noninflammatory stromal cells (fibroblasts and endothelial cells); white: ‘other’ cell class that included nonidentifiable cells, less abundant cells such as macrophages and chondrocytes and ‘normal’ pneumocytes. (A) Correct identification of cancer cells from a mesenchymal tumour (metastatic anaplastic sarcoma) in a snake (GONOXY). (B) A challenging brown-nosed coati (NASNAS) case was diagnosed with a round-cell tumour (lymphosarcoma) where the cancer cell morphology is difficult to be recognised by an algorithm trained with epithelial cells from human lung cancer. (C) A malignant spindle cell tumour from a ring-tailed lemur (LEMCAT) with a haemangiosarcoma disease, as shown, the neoplastic endothelial cells have large rounded nuclei, which may appear morphologically similar to epithelial cancer cells, as opposed to the AI model’s own normal - stromal-endothelial cells. However, the model successfully distinguished the majority of neoplastic from stromal cells. Further complexity is in the occurrence of epithelioid haemangiosarcoma where the cells of origin are endothelial cells but they actually become epithelial-like. (D) In the case of a chimpanzee (PANTRO) with a spindle cell sarcoma, the neoplastic fibroblasts are harder to differentiate from reactive fibroblasts.

In the 18 other vertebrate species (mammalia=11, reptilia=4, aves=2, and amphibia=1), accuracy varies (0.57-0.94). The performance of cancer cells and lymphocyte classification, measured as balanced accuracy, did not vary between tumour types (LR test, cancer cells: median = 0.825, χ^2^[3] = 1.358, p = 0.715; lymphocytes: median = 0.915, χ^2^[3] = 0.308, p = 0.959). However, the classification accuracy of stromal cells differs between tumour types (LR test, median = 0.773,χ^2^[3] = 10.308, p = 0.016), with p-adjusted significant only for differences between epithelial-round cell (z-test, estimate = −0.092, SE = 0.031, z = −3.073, p = 0.018) and mesenchymal-round cell tumour types (estimate = −0.121, SE = 0.039, z = −3.073, p = 0.011). All other comparisons have a p-value higher than 0.05. Surprisingly, in both cases where we reported significant differences, the balanced accuracy of stromal cells in round-cell tumour types was higher than mesenchymal or epithelial tumour types. In our cohort, the round-cell tumour types were present in the dog (*Canis familiaris*), the Tasmanian devil (*Sarcophillus harrisii*), the pygmy goat (*Capra hircus*) and the ring-tailed coati (*Nasua nasua*). These results show a high classification accuracy of the model consistent with expert pathologists’ annotations across tumour types for cancer cells and lymphocytes and slight variations in the case of stromal cells.

### Species and cancer-specific challenges

The detection of cancer cells presented more challenging classifications in lymphosarcoma from the common goat (*Capra hircus*), the ring-tailed coati (*Nasua nasua*) and in lipoma from the dwarf crocodile (*Osteolaemus tetraspis*), which by their cell morphology and tissue architecture may be difficult to be classified by an algorithm trained with epithelial cells from human lung adenocarcinoma (Fig. 4B). These results suggest that the accuracy of computational pathology at single-cell resolution depends on the type of target cancer and its degree of differentiation from the training cancer type. Morphologically complex cancers that drastically change their morphological features or cancers with a high degree of similarity to the normal cells (e.g. lymphosarcoma) represent significant hurdles for transfer learning.

### Morphological preservation across species

To explore the morphological similarity between human and non-human samples, which could explain the accuracy of the AI algorithm, we visualised the morphological space of ~32K cells annotated by expert pathologists using principal component analysis (Fig 5). The PCA analysis was used for dimension reduction (Fig 5) of the 27 features extracted by the AI algorithm at the individual cell level (Table S2). The first three PCA dimensions account for 84.1% of the morphological variance (Fig S3A). The first dimension explains 49.4% of the morphological variance, and the cell features with the highest contributions to that explained variance are associated with nucleus size (area, perimeter, diameter, radius, convex area) and maximum intensity in the colour channels (Fig S3B). These variables are positively correlated with the first dimension, with high importance to explaining individual cells’ position in the morphological space (Fig S3C). The overlap of the volumes in PCA space suggests a high degree of morphological similarity between human and non-human cells (Table S3). For non-human lymphocytes, 84.55% of their morphological space intersects with the human lymphocyte morphospace. And for non-human tumour cells volume, which shows higher morphological variability, 86.49% of its volume is captured by human tumour cells’ volume.

**Figure 5.**
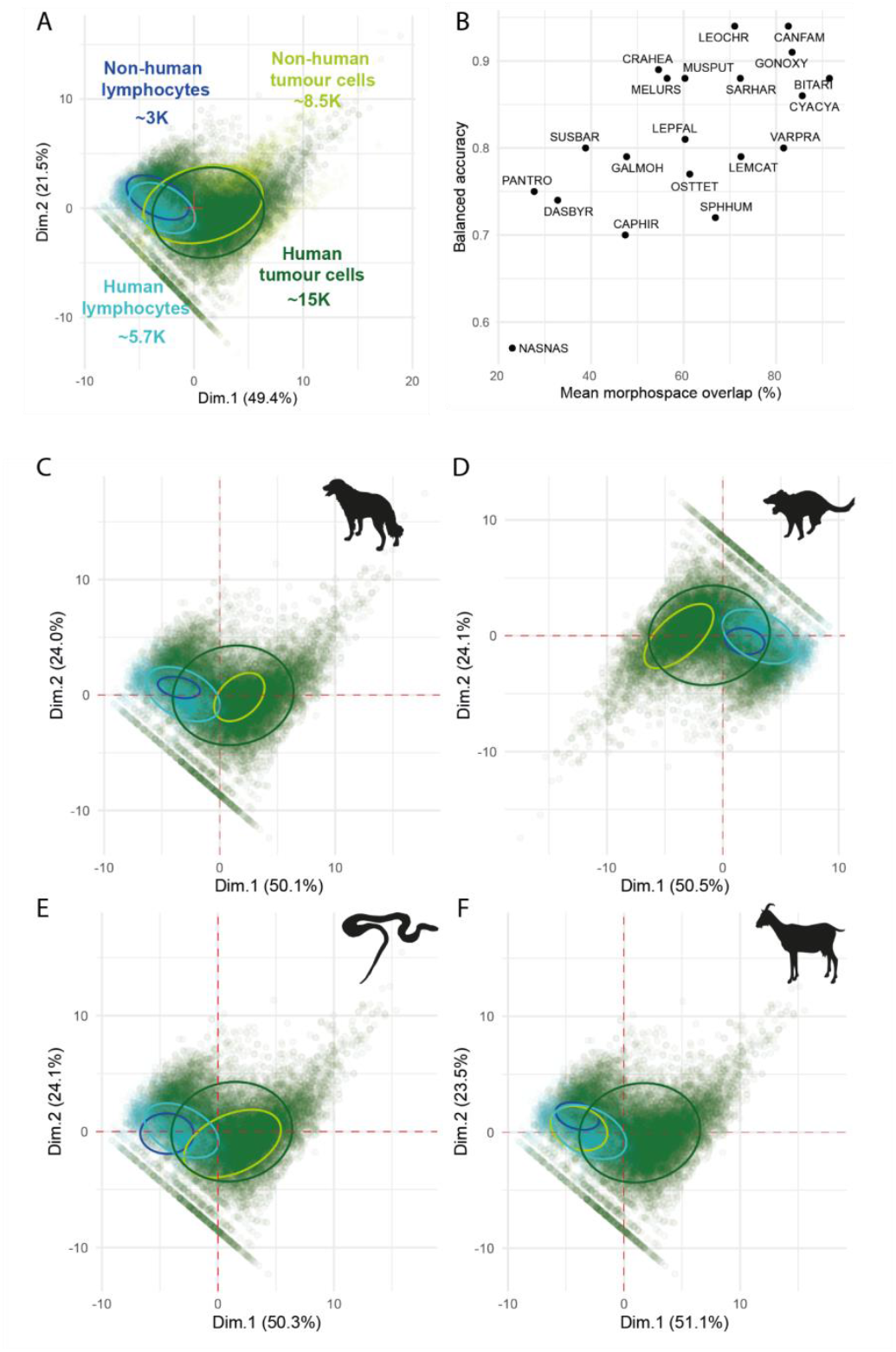
Overlap across the morphological space. (A) Overall high overlap between human and non-human cell morphologies across two dimensions of the principal component analysis, and their explained variances, of the morphological space made by ~31K cells annotated by pathologists. (B) the mean morphospace overlap across animal tumour cells and lymphocytes correlates with the model’s balanced accuracy. (C-F) Species-specific morphological space overlap with human morphospace; (C) *Canis l. familiaris* (CANFAM), (D) *Sarcophilus harrisii* (SARHAR), (E) *Gonyosoma oxycephala* (GONOXY) and (F) *Capra hircus* (CAPHIR). Ellipses denote 95% of the distribution.

### Morphospace overlap as a new guidance metric

To further dissect the relationship between the AI performance and morphological similarity across species, we developed a new metric, termed morphospace overlap, as the average of overlaps of cancer cell/lymphocyte morphological space between a species and humans. We found that the AI model’s balanced accuracy is positively correlated with morphospace overlap (Pearson’s correlation = 0.68, p=0.001; Fig 5B), suggesting that the AI model performed better on species sharing higher morphological similarity with human cells. Species-specific analyses revealed further understanding of the model’s performance. Among the tissues with higher balanced accuracy and high morphospace overlap are dog’s CTVT (Fig 5C), Tasmanian devils’ DFTD (Fig 5D) and snake’s sarcoma (Fig 5E) (morphospace overlap (%) = 82.6, 72.2, and 83.4, balanced accuracy = 0.94, 0.88, and 0.91, respectively) and the goat’s lymphosarcoma (Fig 5F) as one of the challenging cases, with smaller morphological overlap between its tumour cells and human’s cells (morphospace overlap (%) = 47.4, balanced accuracy = 0.7). Species with >70% morphospace overlap had an average of 87.5% balanced accuracy (range 79-94%), and species with >80% morphospace overlap averaged 88% balanced accuracy (range 80-94%). Thus, this new metric may be a useful tool for pathologists to determine the usability of our AI tool.

## Discussion

Comparative oncology pursues the understanding of cancer as a shared phenomenon among species. Here, we have explored the potential of AI through automated pathological image analysis to study cancer morphology and immune response across the tree of life. Previous studies have often been limited to a single species, with applications mainly focused on canine and mouse models (e.g.,^8,13,34^). To the best of our knowledge, this is the first study of computational pathology that includes tumours from vertebrates beyond mammals, such as aves, reptiles and one amphibian. Although the algorithm was trained on human samples, it could distinguish three major cell types with remarkable accuracy in most of the species (19/20 species reached an accuracy ≥ 70% and 12/20 species ≥ 80%). Broadly, our comparative analysis revealed that regardless of species, morphological conservation across species dictates that cells can be detected and correctly classified by a human specimen-trained AI, fostering our endeavour to develop pan-species computational pathology.

Since the model was trained with human epithelial tumour samples, the specimens for testing include other tumour types such as mesenchymal, round cell and neuroendocrine that can have a greater variety of cell morphology, and such diversity of species, and tumour types and sites, likely underpins the wide range of accuracy (0.57-0.94) achieved. For example, in the case of the malignant spindle cell tumour (haemangiosarcoma in a lemur), the neoplastic endothelial cells have large, rounded nuclei, which may appear morphologically similar to that of epithelial cancer cells, as opposed to the elongated nuclei of normal endothelial cells (Fig. 4C). Similarly, for the chimpanzee (*Pan troglodytes*) with a spindle cell sarcoma, the neoplastic fibroblasts are hard to differentiate from reactive fibroblasts with a spindle shape (Fig. 4D). This is a challenge both for the automated analysis and manually by pathologists. Another challenging aspect is the immune compartment, which is highly variable among mammals, birds and reptiles^35^, imposing difficulties that seem complicated to pass with a generic algorithm. Moreover, this is amplified when evaluating cancer affecting the lymphatic tissue, such as lymphosarcoma in the coati (*N. nasua*) and pygmy goat (*C. hircus*), where the white blood cell morphology is altered. Lymphosarcoma cells generally appear similar to normal lymphoid cells, resulting in narrow discriminability chances. In those cases, it may be appropriate to take alternative strategies such as re-train the model, test available models for lymphosarcoma (e.g., ^36^), or develop a new model incorporating other tissue characteristics. To address this issue, we developed a new metric, named morphospace overlap, to guide pathologists who wish to apply the AI tools to their samples based on morphological similarity.

Based on our data, the transferability of existing AI technologies developed for humans to the veterinary domain may be significantly higher than previously thought. Medical treatment for animals has dramatically improved in veterinary clinics, zoological institutions and even wildlife veterinarians^37^, leading to better options for diagnosing and treating cancer in animal patients^23^. Despite these significant advances in veterinary oncology^38^, there are significant constraints and limited availability of veterinary specialists^39^, and consequently, digital tools are not widely used^24,25^. Thus, computational pathology for different species and tumour types will bring tremendous advances for clinical veterinary care and comparative oncology research^24,40^. Many of the advantages are similar to those for human pathology, with the greatest benefits being accessibility to veterinary pathologists, time saved and increased diagnostic accuracy. Significant challenges remain. For instance, our study’s low rate of samples passing quality control highlights a marked difference in sample management between veterinary and human cancer care. Therefore, the pan-species digital pathology atlas, protocols and guidelines for veterinary pathologists provided in this study represent a big step towards rational and efficient transfer of AI technologies to veterinary medicine.

Another potential impact of this study is to empower precision medicine for treating animal cancers. Accurate diagnosis and timely treatment could be critical in preserving endangered and threatened species that represent important breeding populations^41^. We demonstrated how the AI tool can be used to study lymphocytic infiltration in canine transmissible venereal tumours and Tasmanian devil facial tumours with high accuracy and spatial resolution (Fig. 1B-C). As a transmissible disease, the immune response at the organismal level may offer new alternatives to understand the spread of the disease at a population scale from an epidemiological perspective^42,43^. These tumours can colonise a new host by crossing the barriers of histocompatibility associated with the immune system and expressing immunosuppressive cytokines^44,45^. The quantification and spatial detection of both tumour and immune cells can help study immune evasion and treatment in transmissible cancers, building on progress on understanding T cells immune infiltration in Tasmanian devils^46^ and immune regulation in dog’s CTVT tumour regression^47^. Furthermore, a detailed study of the tumour microenvironment can guide new discoveries to understand the mechanisms behind sensitivity and resistance to standard treatments such as chemotherapy,^48,49^. By enabling precision medicine we can advance towards a more personalised and integrative approach to veterinary care^50^.

Comparative oncology also brings tremendous benefits to human cancer research^5,51,52^. Our knowledge of cancer in wild animals is limited, and computational pathology can greatly expand research opportunities that compare cancer in the wild to managed populations, as well as comparisons with human cancer. Cross-species cancer comparisons may help address fundamental questions in cancer biology and evolution. This work revealed highly conserved morphology features across many species, particularly in epithelial and round-cell tumours, highlighting potential evolvability constraints for certain tumour types. The mismatch between species’ evolutionary history and the conserved cellular morphological diversity raises new questions on the origin of cell morphological patterns; is morphological conservation fixed early in metazoan evolutionary history? Or is it the result of stabilising selection imposed by the extracellular matrix to meet homeostatic conditions?^53,54^ Addressing the conserved features and differences in tumour biology can lead to novel research, therapeutics and discoveries that one day could be translated into human and non-human clinical care^37,55^.

Limitations of this study include the limited availability of samples and annotations. It will be important to validate our findings on extended pan-species cohorts and advance our understanding of intratumor heterogeneity across different species and derive more controlled interspecies comparisons. With detailed multiplexing profiles, future attempts can shed more light on immune compositions in the microenvironment.

This work represents a step forward in incorporating machine learning in diagnostic investigations of natural and emerging diseases in animals, enhancing accuracy and sensitivity and complementing veterinary pathologists’ capability in the decision-making process. Computational pathology can bring valuable opportunities for automated diagnosis, tumour grading, scoring, and precision medicine for animal cancers.

## Materials and Methods

In total, 99 H&E samples from 29 species were identified from the Zoological Society of London’s (ZSL) pathological archive, derived from clinical or postmortem examinations of ZSL London Zoo’s living collections (Table S1). Of these, 51 slides from 22 species passed quality control for image analysis, and 18 slides representing 18 species were selected by the pathologists for subsequent analyses. Exclusion criteria were the lack of tumour components and the presence of high amounts of melanin/pigments in the tissue samples hindering the correct identification of individual cells. Samples were either obtained through tissue biopsies from surgery or routine postmortem examinations from animals that were i) examined directly after euthanasia or ii) stored at 4 degrees Celsius and examined within two days of death. A suspect tumour was removed, fixed in 10% buffered formalin solution and trimmed before being sent to external institutions (IZVG Pathology and Finn Pathologists) for histopathological processing, where they were subsequently embedded in paraffin blocks, sectioned and stained with H&E for analysis. Additionally, two samples were provided by the Transmissible Cancer Group, University of Cambridge, as previously reported in the following studies: *Canis familiaris*^56^ and *Sarcophilus harrisii*^9^.

All slides were scanned using NanoZoomer S210 digital slide scanner (C13239-01) and NanoZoomer digital pathology system v.3.1.7 (Hamamatsu) at 40X (228 nm/pixel resolution). The entire deep learning-based single-cell analysis pipeline described in ^33^ was implemented. This pipeline was designed and developed for human lung tumour specimens. Briefly, all 20 whole-section images were first down-scaled to ×20 and then tiled into 2000×2000 images for subsequent three-stage analysis. Firstly, all viable H&E tissue areas are segmented. Secondly, within the segmented tissue image, a spatially-constrained convolutional neural network predicts for each pixel the probability that it belongs to the centre of a nucleus; cell nuclei were then detected from the probability map obtained from the deep network. Lastly, each identified cell was classified using a neighbouring ensemble predictor coupled with a spatially constrained convolutional neural network. There were four cell classes: cancer (malignant epithelial) cells, lymphocytes (including plasma cells), noninflammatory stromal cells (fibroblasts and endothelial cells) and an ‘other’ cell type that included non-identifiable cells, less abundant cells such as macrophages and chondrocytes and ‘normal’ pneumocytes and bronchial epithelial cells.

Because the evaluation of the ‘other’ cell type class would be less mean, given the context of this study, we focused on the three main classes. Two board-certified specialist veterinary pathologists (CP and KH) annotated 14,570 cancer, lymphocyte and stromal single-cell annotations on raw whole-section images.

Features extraction at the cell level was done with two steps: a pre-trained MicroNet model^57^ on lung H&Es to segment all cells, followed by automatic extraction of morphological measurements for the set of properties from each cell’s mask. This allowed the extraction of 27 features for immune and tumour cells annotated by pathologists in the human and non-human slides (MATLAB function ‘regionprops’ with additional modifications as defined in Table S2). Annotated cells were mapped to the segmented cell centroid with a strict threshold of 4 pixels (< 2μm, which is less than 1/3 of a lymphocyte cell), and were visually assessed to confirm correct mapping. Dimension reduction was performed using principal component analysis. Then, we selected the first three dimensions of the PCA, enabling us to build a morphological volume for each cell class. We computed morphological space overlap using the R package ‘dynRB’, which calculates overlap based on the product of overlap at each dimension, the mean overlap across dimensions, or the geometrical mean across the PCA dimensions. We focus on quantifying the percentage of animal cells’ morphological space that is covered by human cells’ morphological space.

The algorithms’ performance for detecting and classifying cells across all species was evaluated directly against the ground truth provided by pathologists’ annotations. Individual class accuracy statistics were calculated using the R function ‘confusionMatrix’ from the R package ‘caret’. To analyse the variability in the classification balanced accuracy values, BCAcc, across tumour or cell types, we fit a generalised linear model considering a beta distribution (logit link function) for continuous values between 0 and 1 (R package betareg). We computed likelihood ratio tests (R package lmtest) to evaluate if the distribution of balances accuracy between tumour types comes from the same χ^2^ distribution. When the χ^2^ test was significant (p < 0.05), we applied multiple comparisons correcting p-values using Tukey’s procedures (R package emmeans). All the statistical tests were performed in R (version 4.0.3) and corresponding R codes are available at https://github.com/simonpcastillo/PanSpeciesHistology.

## Acknowledgements

The authors wish to thank Edmund Flach from the Zoological Society of London, as well as external pathologists Mark Stidworthy, Daniella Denk, Cheryl Sangster, and Ann Pocknell. LMA acknowledges support from the Department of Pediatrics Research Enterprise (University of Utah). Figure icons were taken from phylopic.org (CC BY 3.0), thanks to Sarah Werning, Rebecca Groom, T. Michael Keesey and Tony Hisgett.

## Funding

The Arizona Cancer Evolution Center, University of Arizona, USA.

National Institutes of Health grant U54 CA217376 (YY, AMB, LMA, TAG)

National Institutes of Health grant R01 CA185138 (YY)

Cancer Research UK Career Establishment Award C45982/A21808 (YY)

Cancer Research UK Early Detection Program Award C9203/A28770 (YY)

Cancer Research UK Sarcoma Accelerator C56167/A29363 (YY)

Cancer Research UK Brain tumour Award C25858/A28592 (YY, SPC)

Rosetrees Trust A2714 (YY)

Children’s Cancer and Leukaemia Group CCLGA201906 (YY)

The Royal Marsden Hospital, the ICR National Institute of Health Research Biomedical Research Centre (YY).

Department of Pediatrics Research Enterprise, University of Utah (LMA)

## Author contributions

Conceptualization: KA, SPC, EPM, TAG, CP, YY; Methodology: KA, SPC, CP, KH, HD, SS, EF; Investigation: KA, SPC, CP, YY; Writing: SPC, KA, YY, AMB, LMA, CP, YY, with input from all authors.

## Data and materials availability

The deep-learning pipeline for digital pathology image analysis is previously available for non-commercial research purposes at https://github.com/qalid7/compath. All code used for statistical analyses of image data and morphospace overlap test tool was developed in R (v.4.0.3) and it is available at https://github.com/simonpcastillo/PanSpeciesHistology. A rich, pan-species digital pathology atlas will be made publicly available upon publication, providing pan-species digital slide images, slide digitalisation and quality control protocols, and pathological annotations of 14,570 single-cell annotations across 20 species.

## Supplementary

**Supplementary Figure 1.**
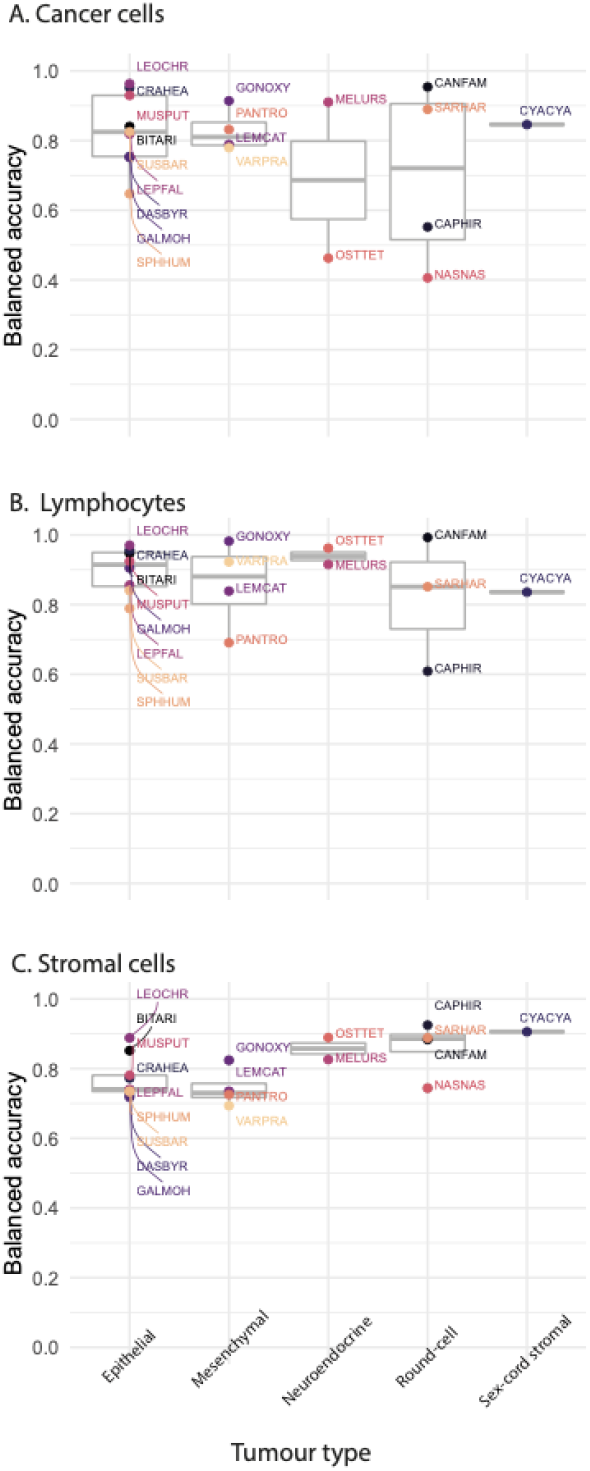
Extended AI single-cell prediction comparison across tumour types. Balanced accuracy is computed as the average of sensitivity and specificity for (A) cancer, (B) stromal and (C) lymphocyte cells for all species. Species are grouped according to their tumour type and are labelled with their codes, for more species information, see Table 1.

**Supplementary Figure 2.**
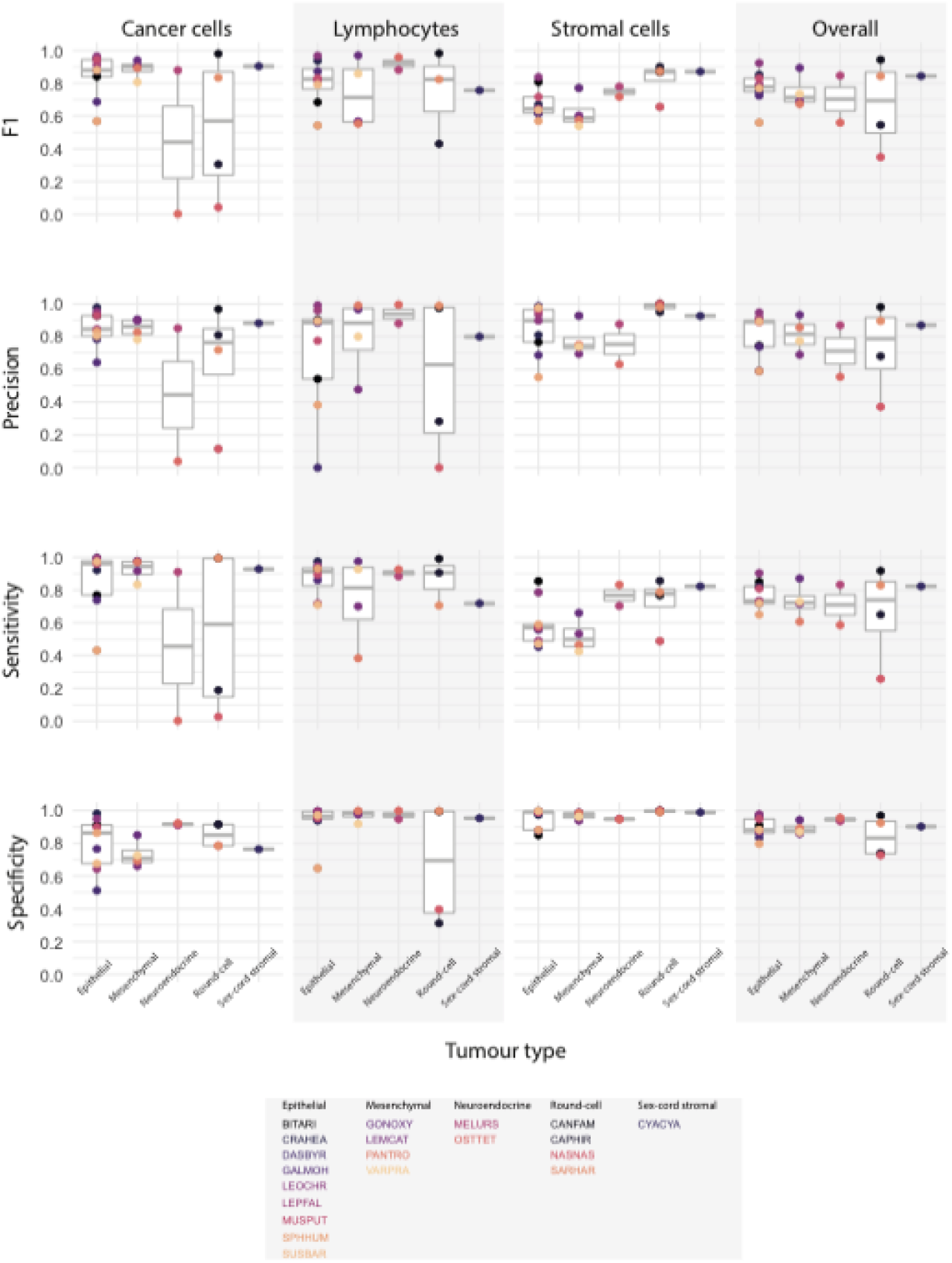
Extended AI prediction variability for inter and intra-species tumour microenvironment cells. For each species, four metrics were evaluated including F1, precision, sensitivity and specificity (rows) for the prediction accuracy of cancer, lymphocyte and stromal cells as well as their average shown as ‘overall’ (columns). Species are grouped according to their tumour type and are labelled with their codes, for more species information, see Table 1.

**Supplementary Figure 3.**
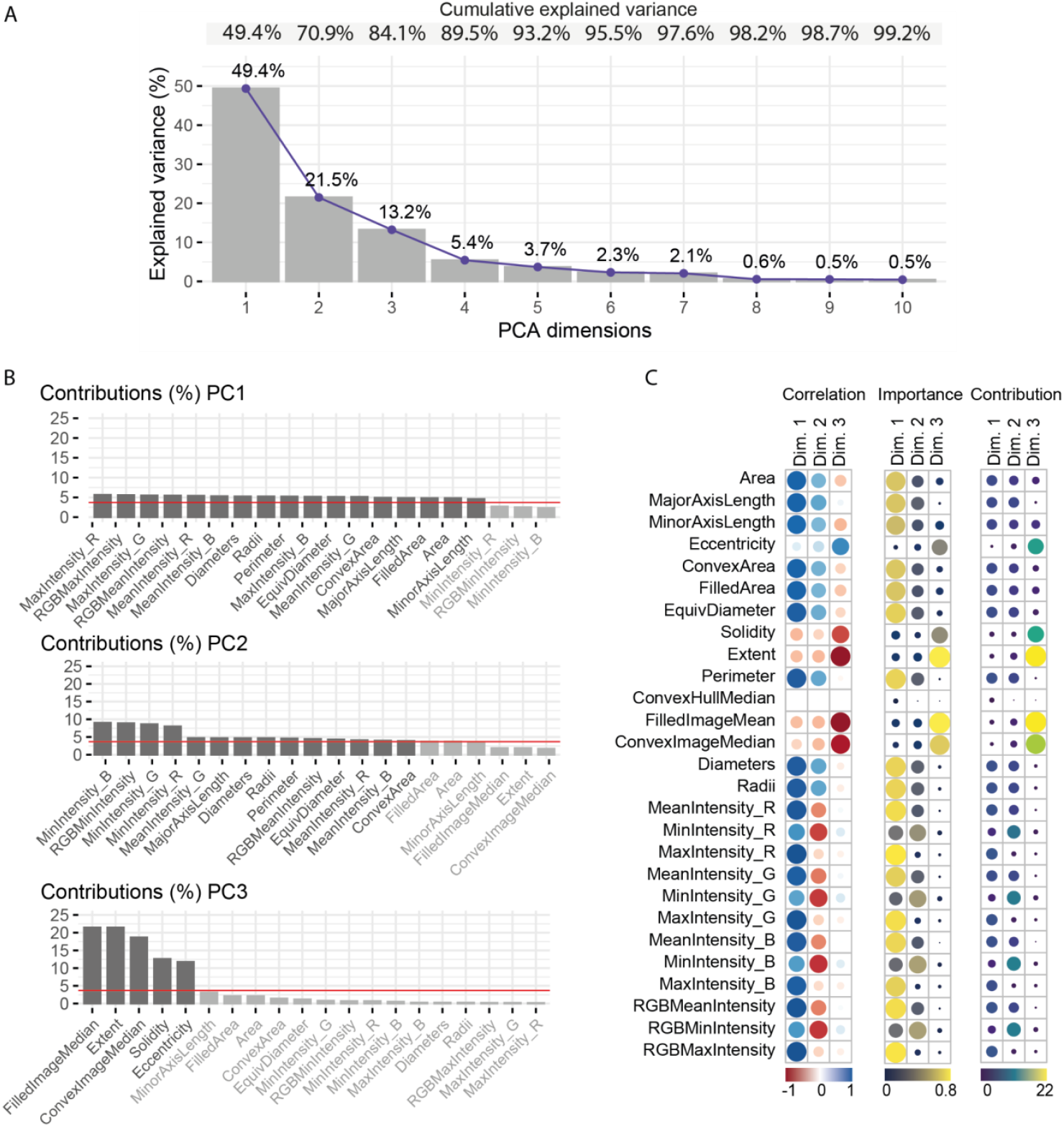
Analysis of the morphological space. (A) Dimensions of the principal component analysis (PCA) and their explained variances. (B) The highest 20 contributions to PCA dimensions’ explained variances. Darker bars are features above the mean contribution (red line). (C) From left to right, correlation, importance and contribution of the single-cell morphological features to PCA dimension.

**Supplementary Table 1.** Summary of sample preparation methods as provided from the Zoological Society of London’s pathological archive.

| Case ID | Species code | Pathologists | Method |
| --- | --- | --- | --- |
| **B01/17** | MUSPUT | IZVG/RVC- DD | Biopsy: removed during surgery and formalin-fixed |
| **B02/18** | GALMOH | IZVG/RVC- DD | Biopsy: removed during surgery and formalin-fixed |
| **B04/17** | LEMCAT | IZVG/RVC- MS | Biopsy: removed during surgery and formalin-fixed |
| **B07-8/04** | PANTRO | ZSL- AP | Biopsy - removed during surgery and formalin-fixed |
| **B09/04** | DASBYR | ZSL- AP | Biopsy - removed during surgery and formalin-fixed |
| **W17M035** | MELURS | IZVG/RVC- MS | Euthanasia: Carcass fresh – PM examination one day after death |
| **W17R187** | OSTTET | IZVG/RVC- DD | Natural death: Carcass fresh - PM on day of death |
| **ZA1360/15** | LEPFAL | IZVG/RVC- MS | Natural death: Carcass slightly autolysed – PM on day of death |
| **ZB017/18** | CYACYA | IZVG/RVC- MS | Euthanasia: Carcass fresh – PM examination one day after death |
| **ZB485/19** | SPHHUM | IZVG/RVC- CS | Euthanasia: No comment on carcass condition - PM carried out 2 days after euthanasia |
| **ZM134/17** | CAPHIR | IZVG/RVC- CS | Carcass fresh – euthanised and PM'd on day of death |
| **ZM138/17** | SUSBAR | IZVG/RVC- MS | Carcass fresh - PM on day of death |
| **ZM203/17** | LEOCHR | IZVG/RVC- MS | Carcass fresh - PM on day of death |
| **ZM633/18** | CRAHEA | IZVG/RVC- DD | Euthanasia: Carcass fresh – PM examination one day after death |
| **ZM748/18** | NASNAS | IZVG/RVC- CS | Euthanasia: Carcass fresh – PM examination one day after death |
| **ZR1145/15** | GONOXY | IZVG/RVC- MS | Euthanasia: Carcass fresh - kept in fridge two days before examination |
| **ZR1148/18** | VARPRA | IZVG/RVC- DD | Euthanasia: Carcass fresh – PM examination one day after death |
| **ZR474/19** | BITARI | IZVG/RVC- CS | Euthanasia: Carcass fresh – PM examination one day after death |

**Supplementary Table 2.** The 27 single-cell features extracted to compute the morphological space.

| Feature | Description |
| --- | --- |
| Area | Two-dimensional extension of a shape |
| MajorAxisLength | Longest diameter |
| MinorAxisLength | Shortest diameter |
| Eccentricity | Magnitude inversely related to shape curvature |
| ConvexArea | Area resulting from connecting the external points of the shape |
| FilledArea | Area of a corresponding image with holes filled in |
| EquivDiameter | Diameter of a circle with the same area as the region |
| Solidity | Extent to which the shape fills the convex area |
| Extent | Ratio of pixels in the region to pixels in the total bounding box |
| Perimeter | Length of the shape boundary |
| ConvexHullMean | Smallest convex polygon that can contain the region |
| FilledImageMean | Average of pixels corresponding to the segmented mask, with all holes filled |
| ConvexImageMean | Average of pixels corresponding to a segmented mask which specifies the convex hull of the region |
| Diameters | Cell diameter using major and minor axes |
| Radii | Cell radius |
| MeanIntensity_R | Mean pixel intensity in the red channel |
| MinIntensity_R | Minimum pixel intensity in the red channel |
| MaxIntensity_R | Maximum pixel intensity in the red channel |
| MeanIntensity_G | Mean pixel intensity in the green channel |
| MinIntensity_G | Minimum pixel intensity in the green channel |
| MaxIntensity_G | Maximum pixel intensity in the green channel |
| MeanIntensity_B | Mean pixel intensity in the blue channel |
| MinIntensity_B | Minimum pixel intensity in the blue channel |
| MaxIntensity_B | Maximum pixel intensity in the blue channel |
| RGBMeanIntensity | Mean pixel intensity in the composed image |
| RGBMinIntensity | Minimum pixel intensity in the composed image |
| RGBMaxIntensity | Maximum pixel intensity in the composed image |

**Supplementary table 3.** Morphological volumes overlap of human cells on non-human cells’ morphological space calculated by the three methods. The highest overlap values for non-human lymphocytes and tumour cells are bold-faced.

| Morphological volume of | Covered by the volume of | % Overlap (product) | % Overlap (mean) | % Overlap (geom. mean) |
| --- | --- | --- | --- | --- |
| **Non-human lymphocytes** | **Human lymphocytes** | **63.47** | **84.55** | **81.25** |
| Non-human lymphocytes | Human tumour cells | 40.41 | 67.6 | 59.84 |
| Non-human tumour cells | Human lymphocytes | 16.56 | 54.82 | 50.24 |
| **Non-human tumour cells** | **Human tumour cells** | **67.97** | **86.49** | **86.14** |

